# Structured environments fundamentally alter dynamics and stability of ecological communities

**DOI:** 10.1101/366559

**Authors:** Nick Vallespir, Lowery & Tristan Ursell

## Abstract

The dynamics and stability of ecological communities are intimately linked with the specific interactions – like cooperation or predation – between constituent species. In microbial communities, like those found in soils or the mammalian gut, physical anisotropies produced by fluid flow and chemical gradients impact community structure and ecological dynamics, even in structurally isotropic environments. Though natural communities existing in physically unstructured environments is rare, the role of environmental structure in determining community dynamics and stability remains poorly studied. To address this gap, we used modified Lotka-Volterra simulations of competitive microbial communities to characterize the effects of surface structure on community dynamics. We find that environmental structure has profound effects on communities, in a manner dependent on the specific pattern of interactions between community members. For two mutually competing species, eventual extinction of one competitor is effectively guaranteed in isotropic environments. However, addition of environmental structure enables long-term coexistence of both species via local ‘pinning’ of competition interfaces, even when one species has a significant competitive advantage. In contrast, while three species competing in an intransitive loop (as in a game of rock-paper-scissors) coexist stably in isotropic environments, structural anisotropy disrupts the spatial patterns on which coexistence depends, causing chaotic population fluctuations and subsequent extinction cascades. These results indicate that the stability of microbial communities strongly depends on the structural environment in which they reside. Therefore, a more complete ecological understanding, including effective manipulation and interventions in natural communities of interest, must account for the physical structure of the environment.

**SIGNIFICANCE:** Many microbial communities of ecological and medical importance reside in complex and heterogeneous environments, such as soils or intestinal tracts. While many studies consider the effects of flow or chemical gradients in structuring these communities, how the physical structure of the environment shapes community dynamics and outcomes remains poorly understood. Using simulations of competitive microbial communities, we show that stability and dynamics qualitatively shift in environments with complex surface structures compared to open isotropic environments. Therefore, in addition to biochemical interactions between species, our work suggests that the physical structure of the environment is an equally important determinant of dynamics and stability in microbial communities, in a manner dependent on the specific patterns of interactions within that community.

## INTRODUCTION

From the scale of large metazoans down to microbes, natural environments are replete with multi-species communities that compete for resources and space, and in many cases actively predate other species within their environment. Within complex ecosystems the topology and type of interactions between constituent species are thought to be a primary determinants of ecosystem dynamics and stability. Typical pairwise interactions, like competition, cooperation, or predation, form the building blocks for constructing multi-species interactions and can be used to predict dynamics and stability in ‘well-mixed’ environments where spatial distributions are uniform (1, 2). Interaction topology plays a particularly important role in species coexistence. For instance, in three-species intransitive competition (as in the classic rock-paper-scissors game), extinction of any species results in extinction cascades that favor dominance of a single species. Microbial systems present a particularly salient manifestation of these concepts, not only because complex communities of microbes are found in a wide array of industrial- and health-relevant environments, like soils and the mammalian gut, but also because the ability to genetically recapitulate and manipulate specific pairwise interactions biochemically makes microbial systems particularly well-suited for testing our understanding of fundamental mechanisms underlying ecosystem dynamics.

Characterization of interactions within ecological networks, and their corresponding biochemical mechanisms, often focuses on microbial communities in which the spatial distribution of actors can significantly impact the type and magnitude of those interactions, and the resulting population dynamics. For example, spatially localized clonal domains that result from competition between mutually killing isolates of *Vibrio cholerae* may facilitate emergence of cooperative behaviors like public good secretion (3). Similarly, large clonal domains stabilized three-way intransitive competition within a consortium of *E. coli* strains (4); the same consortium was unstable in well-mixed environments. Reversing the causative arrow, ecological interactions can also dictate spatial arrangements of genotypes: in simulated three-species intransitive consortia with mobile individuals, lack of a single dominant competitor leads to population waves that continually migrate throughout the environment (5), thereby ensuring dynamic and long-term stability in species representation. Conversely, in competition between two mutual killers, coarsening of clonal domains guarantees the eventual extinction of one of the species (3), unless additional interaction mechanisms are present (6). Therefore, in contrast to dynamics that play out in well-mixed environments, it is clear that the spatial distribution of organisms is an important determinant of community dynamics and long-term ecological outcomes.

A common condition imposed on simulations of spatially explicit ecological systems is environmental isotropy – defined by the system having the same chemical and physical properties in all directions (for example, a homogeneous 2D plane (3, 5, 7)). While such simplifications are essential in building fundamental understanding of system dynamics, they do not reflect salient environmental anisotropies found in most natural systems, such as chemical gradients, fluid flow, and complex surfaces. Despite relevance to natural communities, examination of the mechanisms by which environmental anisotropies affect ecological communities is sparse. In single species populations, colonizing complex environments can result in drastic changes in spatial distributions. For instance, using microfluidic devices, Drescher *et al*. showed that surface morphology and fluid shear forces interact to drive formation of novel biofilm structures in *Pseudomonas aeruginosa* (8). Biofilm formation can also disrupt fluid flow in a microfluidic mimic of soil environments, which in turn allows for coexistence of competing cheater and cooperator phenotypes of *P. aeruginosa* that are otherwise unstable under well-mixed conditions (9). Importantly, these perturbations to population structure are commensurate with length scales at which mixing occurs for *in vivo* communities such as the mammalian (10, 11) and fish (12) guts, or in dental plaque (13). Theoretical investigations indicate that similar environmental perturbations are likely to affect multispecies communities: for example, turbulent flow can disrupt spatial patterning of intransitive three-species communities and thus increase the risk of extinction cascades (14), while graph theoretic approaches suggest that random perturbations to spatial lattices result in similar community destabilization (15). Together, these results suggest not only that spatial distributions of organisms influence ecological dynamics, but that the magnitude of these effects depends strongly on the specific nature of anisotropies within the environment.

In this work, we systematically characterized the effects of structural anisotropy on multi-species population dynamics and spatial distributions within *in silico* ecological communities. The structural attributes of these simulations are intended to capture the primary spatial structure found in natural environments, like the packing of steric soil particles or the contents and epithelial structure of the mammalian gut. Using reaction-diffusion models, we simulated asymmetrically competing two-species and intransitively competing three-species ecological networks in the presence of steric barriers arranged in a lattice within the environment. These networks and the corresponding simulations were chosen for direct comparison to previous work (3, 5) which provide clear expectations for spatial distributions and community dynamics in homogeneous environments, and which we discuss in context below. We find that the addition of environmental structure fundamentally alters community dynamics in both two- and three-species competitive systems. In the two-species case, coarsening of genetic domains that would otherwise lead to extinction of one competitor is arrested due to ‘pinning’ of competition interfaces between barriers, resulting in long-term coexistence of both species. This effect is strongly linked to the geometry of the steric barriers, and is robust to asymmetry in competitive fitness. For intransitive three-species competition, steric barriers cause interference between traveling population waves, inducing chaotic fluctuations in the abundances and spatial distributions of species and a concomitant increase in the probability of extinction cascades. Our results affirm that the trajectories, stability, and spatial structure of ecological communities are drastically altered by the structure and length scale of structural perturbations in the environment.

## RESULTS

### Competition model

We model interspecies interactions using an adapted version of the Lotka-Volterra (LV) competition framework. In the classic LV model, interaction mechanisms and fitness differences are combined into a single parameter, which realizes competition as a reduced effective carrying capacity for the focal species relative to the density of a competitor – hence there is no differentiation between e.g. competition for space and toxin-mediated killing. Here, we extend the classic framework to reflect ‘active competition’, where passive competition for space and nutrients (affecting carrying capacity) is decoupled from active competition mechanisms that directly impact growth rate, such as T6SS mediated killing or bacteriocin production (16, 17), giving the partial differential equation (PDE)

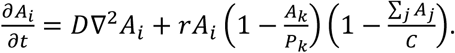

Here *A_i_*, is the local concentration of focal species *i, A_k_* is the active competitor species for *A_i_*, and the sum is over all species passively competing for space and nutrients. The primary dispersal mechanism is through diffusion characterized by *D*, basal growth rate is given by *r*, and carrying capacity by *C*. Active competition is characterized by the concentration parameter *P*, where lower values of *P* indicate more potent active competition (i.e. lower concentrations of the active competitor are required to cause death). This framework explicitly models passive fitness differences (through *C)* and anti-competitor mechanisms (through *P)*, thereby capturing two basal and distinct mechanisms of microbial competition. This model is appropriate for describing local competitive interactions, like contact-mediated killing or local killing by secreted toxins. Additional PDEs would be required to describe highly motile cells, exogenous gradients, or the production, potency, and transport of rapidly diffusing secreted toxins. This set of PDEs establishes a baseline set of assumptions and corresponding phenomena from which to build more complex models.

In this work, we focus specifically on the competitive effects, and assume constant growth rates *r*, diffusion *D*, and carrying capacities *C* for all species in the community. This simplification allows the population density to be scaled by carrying capacity and the time to be scaled by the growth rate, which reduces the parameter space of the model leaving the dimensionless version of *P* (i.e. *P/C*) as the single free parameter that dictates the strength of active interspecies competition

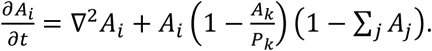

Here time is in units of *r*^-1^, length is in units of 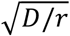, and organismal concentrations *A_i_*· (i.e. number per unit area) are in units of *C*, therefore 0 ≤ *A_i_* ≤1. The natural length scale 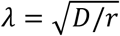 is proportional to the root mean squared distance an organism will move over a single doubling time.

We used this non-dimensionalized model to simulate communities in a 2D environment into which we introduce structural anisotropy via a lattice of steric pillars (see Figures 1 and 3). Like a grain in soil or tissue in a gut, these pillars do not allow free transport through them, nor microbes to occupy them; their perimeter is a reflecting boundary condition. Structural perturbations were explored by introducing a triangular lattice of steric circular pillars, with each lattice fully characterized by the radii of the pillars *R* and the center-to-center spacing of the pillars Δ*x*, with each simulation evolving in a square domain of side length *L*. These parameters (pillar radius *R*, pillar spacing Δ*x*, and simulation size *L*) are reported in units of *λ*. We then characterized the impact of these perturbations on the spatial distribution and dynamics of *in silico* communities across structural length scales by monitoring the distributions and abundances of resident community members as we varied the radius and density of pillars within the simulation environment.

**Figure 1:**
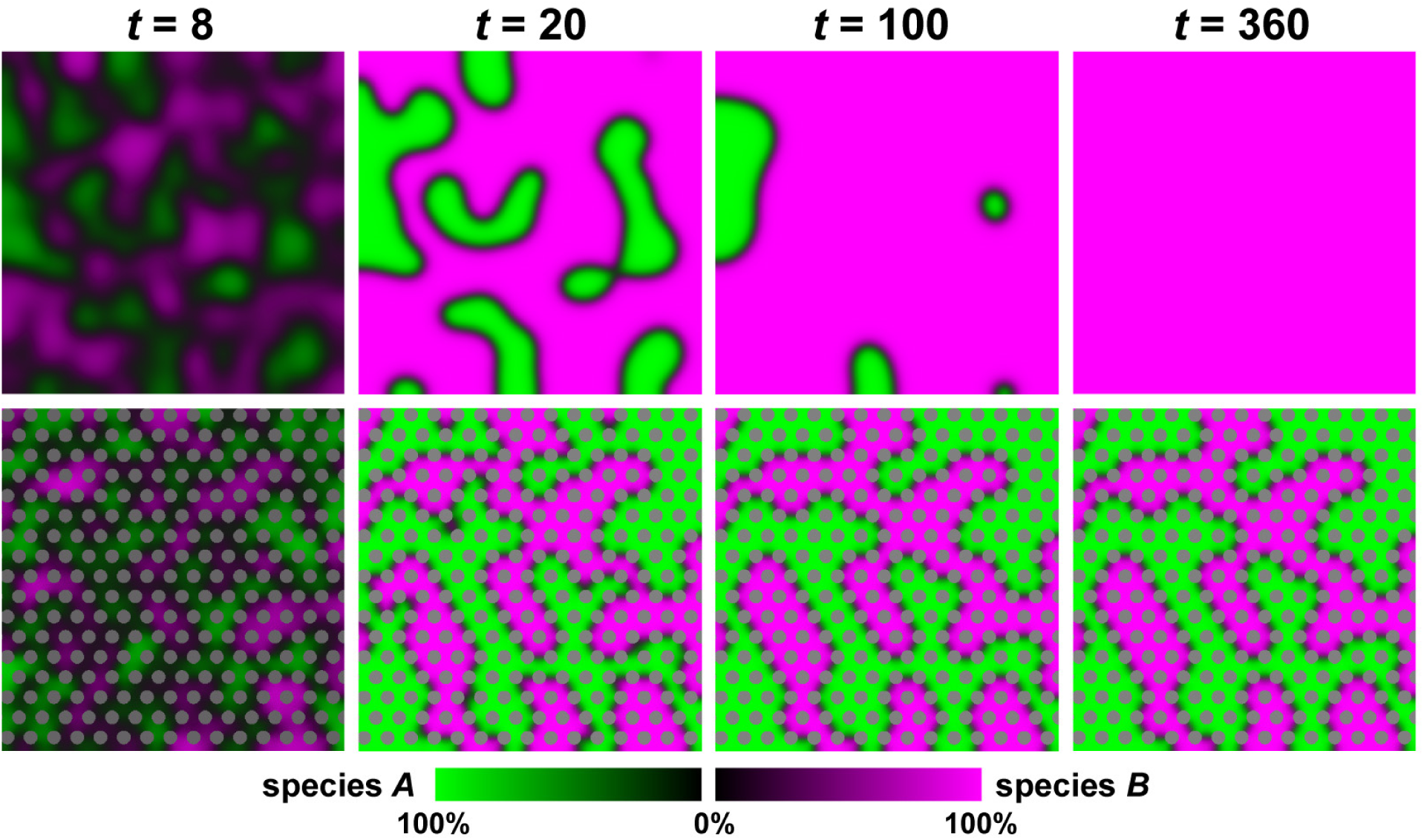
Structurally anisotropic environments arrest genetic phase separation in two-species systems, resulting in long term coexistence. Panels depict snapshots from simulations of two-species competition in structurally isotropic (top row) and anisotropic (bottom row) environments, with color intensity reflecting species abundance, and pillars shown in grey. Time is measured in doubling times. Under isotropic conditions, domain coarsening robustly leads to extinction of one of the species. Anisotropic environments, however, allow for local pinning of competition interfaces, resulting in arrest of domain coarsening and thereby sustained coexistence. Simulation parameters are *L* / (1.29 *λ*) = 100 and *P* = 0.1, with *R* / (1.29 *λ*) = 2 and Δ*x* = 3.4 *R* for the anisotropic case.

**Figure 2:**
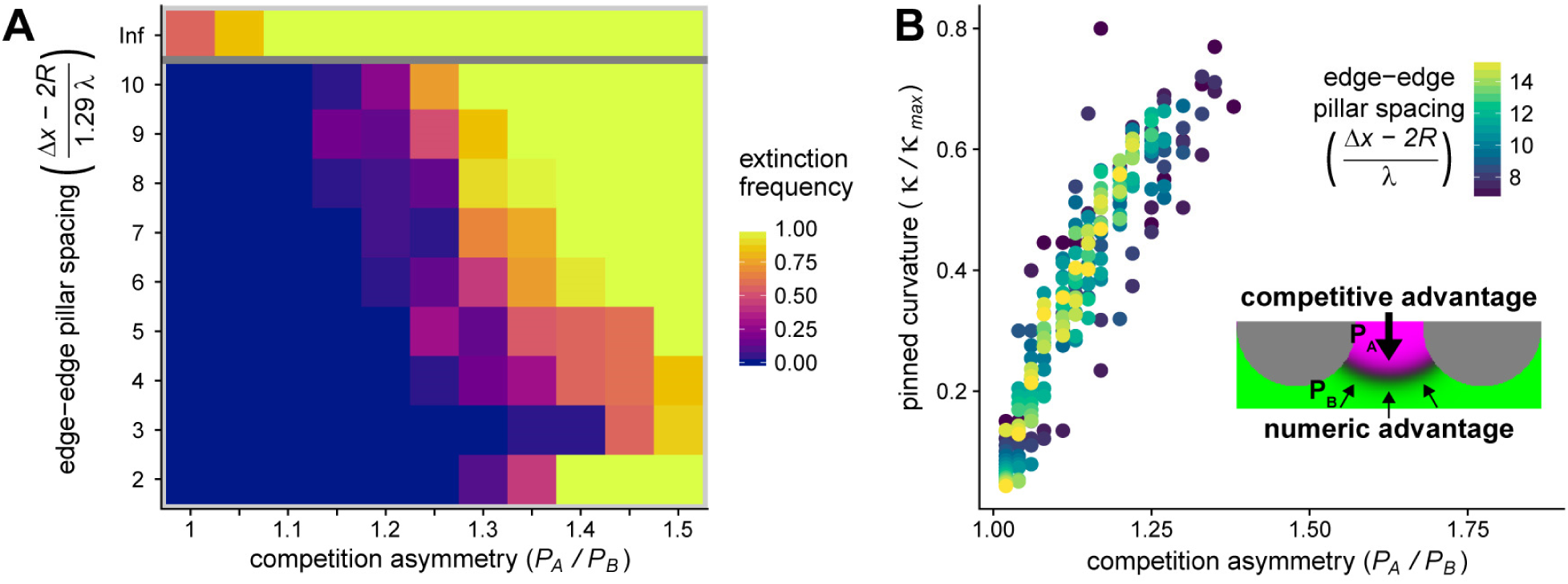
Coexistence of species with asymmetric competitive fitness is maintained by pinning of competition interfaces. **A**, Extinction frequency from 30 replicate simulations per coordinate over 2000 doubling times as a function of competition asymmetry and pillar spacing. Here, the ratio Δ*x* / *R* is held constant at 3, while varying the lattice constant Δ*x* relative to the natural length scale *λ*. Data for isotropic environmental conditions (no pillars) are depicted above the grey line – note that the reason some simulations were not observed to go extinct was due to insufficient simulation duration; with more time, all isotropic simulations would go extinct. Higher pillar density stabilizes coexistence between strains with larger competitive asymmetries, up to the point at which the environment cannot sustain sufficiently large domains to stabilize the competition interface; this produces the increase in extinction frequency at bottom right (see Supplemental Movie 2). Simulation parameters are *L* / (1.29++ *λ*) = 100 and the average competition strength (*P_A_* + *P_B_*)/2 is held constant at 0.1. B, Stable interface curvature (*k*) relative to the maximum possible interface curvature (*k*_max_) as a function of competitive asymmetry (see Supplemental Text 1). Edge-to-edge pillar spacing is indicated by point color. Pinning and stable coexistence was observed for competitive asymmetries greater than 1.35 (see panel A), but were omitted in B because pillar spacings were too small for reliable curvature estimation. The inset schematically depicts stable curved interfaces, where the numerical advantage of the weaker competitor species (green) balances the advantage of the stronger competitor species (magenta). See methods for description of curvature calculation.

**Figure 3:**
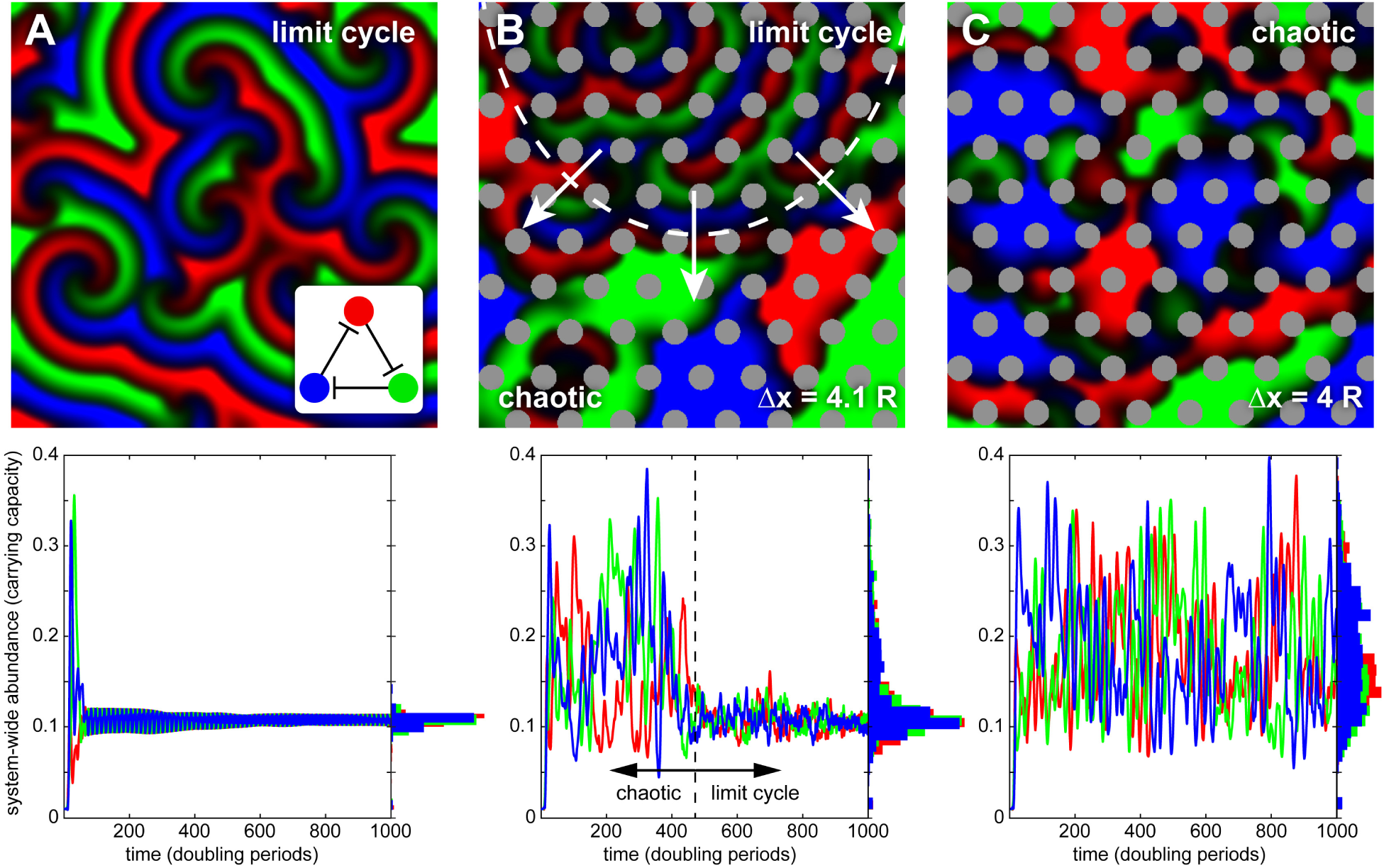
Structurally anisotropic environments disrupt spatial patterns and cyclic dynamics in intransitively competing 3-species communities. Top panels show snapshots of spatial distributions from representative simulations, with corresponding abundance dynamics plotted below. A, following a brief ‘grow-in’ period, isotropic conditions result in spiral waves and cyclic abundance dynamics with corresponding stable coexistence. B, introduction of pillars disrupts cyclic pattern formation, leading to irregular spatial distributions and large fluctuations in species abundance. However, in this example the system eventually transitions into a stable cyclic state, indicated by dashed boundaries in top and bottom panels, with arrows in the top panel indicating the direction of the expanding cyclic region. C, more densely packed pillars hinder transition to a limit cycle, resulting in sustained large fluctuations in abundance and irregular species distributions. Simulation parameters are *L* / *λ* = 158, *P* = 0.1, and for simulations including pillars *R* / *λ* = 4.74 and Δ*x* as indicated.

### Competition between two mutual killers

#### Structured environments arrest genetic phase separation

For an actively competing two-species community in an isotropic environment, recent theoretical and experimental work indicates that species phase separate according to genotype, with the eventual extinction of one species via domain coarsening (3). In contrast, we find that when morphological structure is introduced into the environment genetic phase separation is arrested, resulting in stable coexistence of mutually killing genotypes (Figure 1, Supplemental Movie 1). Arrest occurs by ‘pinning’ of competition interfaces between steric barriers (i.e. pillars). In both isotropic and anisotropic environments, coarsening of genetic domains is driven by the curvature of competition interfaces. If competition is symmetric, a flat interface will not move, whereas a curved interface will translate toward the center of the circumscribing circle. In isotropic conditions, stable flat interfaces are the exception, only found in the rare case where a single flat interface bisects the entire environment, which is itself increasingly unlikely in larger environments. Thus, all domains enclosed by a competitor will eventually be consumed and one of the competitors will go extinct. In contrast, we find that flat competition interfaces are stabilized between steric barriers, resulting in the arrest of domain coarsening and subsequent long-term coexistence of both species (Figure 1). Importantly, for symmetric competition we observed that the size and/or density of pillars had little effect on community stabilization (left edge of Figure 2A), suggesting that for well-matched competitors even slight structural perturbations that allow for interface pinning may be sufficient to foster coexistence.

#### Pinning of genetic domain interfaces is robust to asymmetric competition

When one species is a more potent competitor (e.g. *P_A_* > *P_B_*), even the symmetry of an environment fully bisected by a linear competition interface will result in extinction of the weaker competitor. While flat interfaces balance symmetric competition, they are not stable when one species has a competitive advantage, and instead will translate through space. Likewise, when competition is asymmetric in an isotropic environment, over an ensemble of random initial conditions the dominant competitor will drive the weaker competitor to extinction in the overwhelming majority of cases. We wanted to know if structural perturbations could stabilize coexistence even when competition was asymmetric. Thus we performed simulations identical to those described above, but varied the ratio of the competition parameters, *P_A_/P_B_*, while holding their mean constant. We observed that stable coexistence via interface pinning was robust to asymmetric competition (Figure 2, Supplemental Movie 2) within certain regimes of the lattice parameters. The mechanism, however, was somewhat counterintuitive: for a given degree of competition asymmetry, *P_A_/P_B_*, there exists some critical interface curvature that balances the numeric advantage of the weaker species against the competitive advantage of the more potent species (Figure 2B, inset). This is true regardless of the presence of environmental structure; however, in isotropic conditions this competitive equilibrium is unstable, and any perturbation of domain curvature will result in interface translation and eventual extinction. We found that structural perturbations stabilize the competitive equilibrium created by curved competitive interfaces if the spatial structure of the environment can support the critical curvature between two steric surfaces (Supplemental Text 1) -- only then will phase separation halt and coexistence be maintained. Otherwise, the dominant competitor will drive the weaker species to extinction (Figure 2A), with slower dynamics than isotropic conditions.

Using geometric and scaling arguments (Supplemental Text 1), we predicted that the critical curvature should be an approximately linear function of the competitive asymmetry and confirmed this with our simulations (Figure 2B). Unlike symmetric competition, where coexistence is fully determined by flat competition interfaces, the curved interfaces required to equilibrate asymmetric competition also impose a minimum stable domain size on the competitively disadvantaged species that depends on the lattice parameters. This is because a sufficiently large population of weak competitors is required to compensate for competitive losses at the interface through growth and diffusion (note the increased levels of extinction with the smallest pillar spacings in Figure 2A, and the dissolution of domains in Supplemental Movie 2 that were stable under the symmetric competition of Supplemental Movie 1).

### Three species intransitive competition

#### Environmental structure disrupts three-species dynamics

Previous *in silico* simulations of an intransitively competing three-species network (i.e. displaying a cyclic competitive hierarchy, as in the game rock-paper-scissors) within an isotropic environment resulted in the formation of striking spiral wave patterns, in which dense waves of species constantly migrate throughout the environment, with each species wave chasing its prey and being followed by its predator (see (5), and recapitulated in our model in Figure 3A). Despite constant flux of species at small length scales, the community exhibited stable coexistence of all three species on ecological time scales (more than 10^4^ generations) when provided with a sufficiently large environment relative to the natural length scale set by diffusion and growth. These findings agree with the earlier experimental results of Kerr et al. (4), albeit at different time and length scales. However, it should be noted that previous theoretical work indicates that fluctuations (18) or finite number effects (19) can force such systems into heteroclinic cycles that eventually lead to extinction cascades.

Given the drastic changes in ecological outcomes when structural perturbations were introduced in two species competitive systems, we wanted to characterize how dynamics and outcomes changed in three species competition when we included structural perturbations. We performed simulations using the same set of governing equations as in the two species case, now accounting for the topology a cyclic competitive hierarchy and imposing fully symmetric competition for simplicity. We found that the introduction of spatial structure into the environment significantly destabilizes wave patterns observed under isotropic conditions in a manner that strongly depends on the spacing and size of steric barriers (Figure 3). For example, while densely packed barriers prevent regular pattern formation and result in erratic fluctuations in species abundance (Figure 3C), increasing the space between pillars by a small amount allows the system to re-establish wave patterns that dominate the environment and significantly reduce the magnitude of population fluctuations (Figure 3B). We therefore set out to characterize the complex dynamics arising from intransitive competition in structured environments, with special attention paid to transitions in population dynamics as a function of quantitative changes in environmental structure.

#### Introducing structural anisotropy leads to chaotic fluctuations in species abundance and extinction cascades

To quantify how structural perturbations destabilize pattern formation and cyclic dynamics in our deterministic simulations, we examined the dynamic trajectories of multiple replicates of the same steric pillar array initialized with controlled, random differences in the initial distributions of the three species. We then compared the correlations in species distributions between replicate simulations as the system evolved. In contrast to limit-cycle dynamics in isotropic environments, we found that increasing pillar density resulted in extreme sensitivity to perturbations of initial conditions with an exponential decay in initial correlations through time (Figure 4), a hallmark of chaotic dynamics (20). Chaotic fluctuations were accompanied by rapid transitions into extinction cascades (evident in Figure 4, where correlation traces are truncated at the first extinction event among replicates).

**Figure 4:**
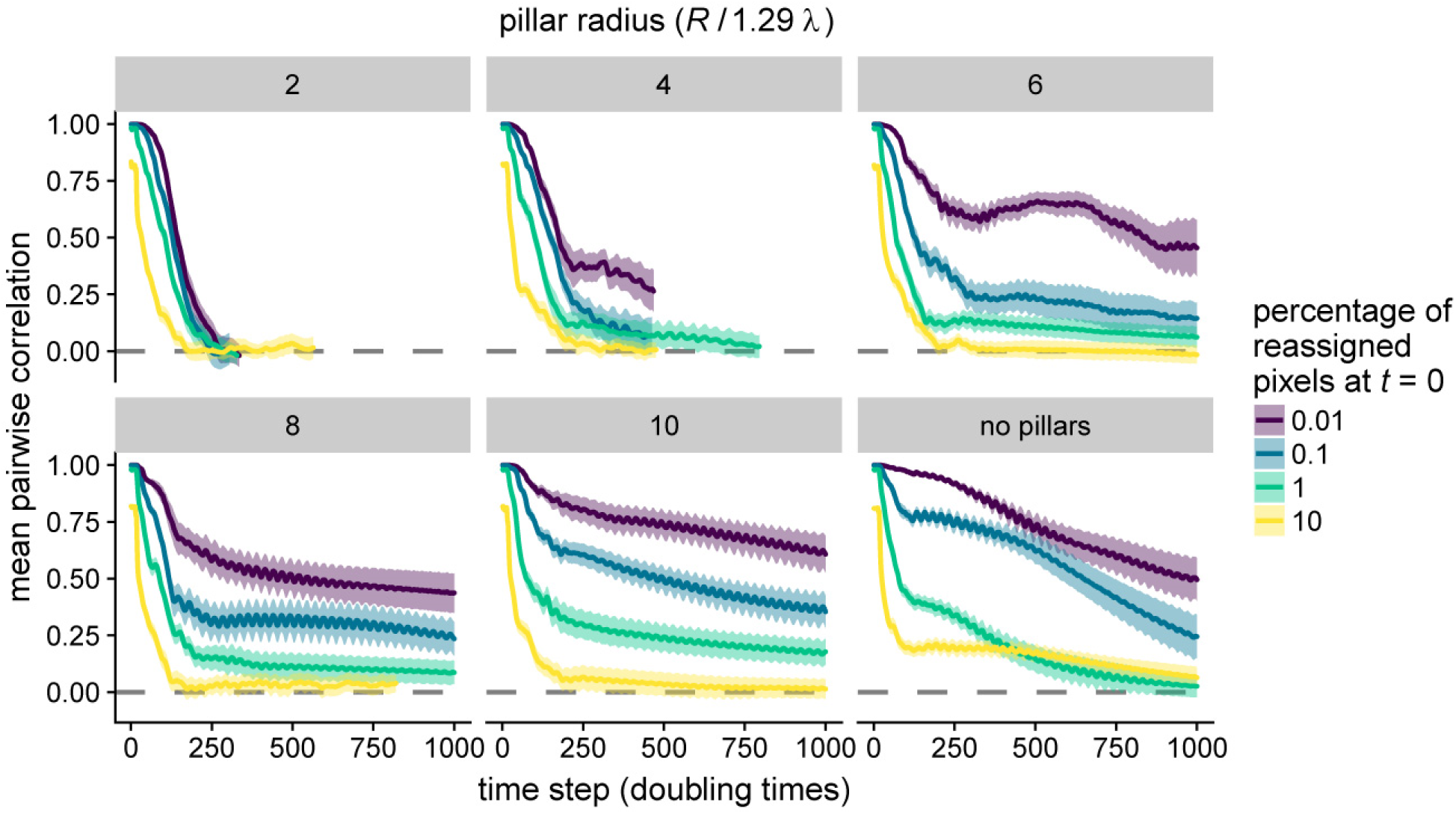
Ecological dynamics display extreme sensitivity to initial conditions depending on environmental structure. In each panel, 10 replicate simulations were identically initialized, then a small percentage (indicated by line color) of grid locations in each replicate were randomly re-sampled creating correlated initial conditions. Each panel shows the spatial (pixel-by-pixel) correlation over time between replicate simulations, averaged over all 45 unique pairwise comparisons. Shaded regions indicate standard error of the mean. As the pillar array becomes denser, rapid decorrelation among replicates results from minute perturbations to initial conditions, a hallmark of chaotic dynamics. In contrast, as pillar spacing increases, some fraction of simulations fall into a limit cycle and thus have non-zero correlations with similar initial conditions. Simulation parameters are *L* / (1.29 *λ*) = 100, *P* = 0.1, and Δ*x* = 4 *R*, with pillar size *R* indicated at the top of each panel.

In initial simulations, we noted that species distributions often exhibited dynamic transitions between patterns of spiral waves and chaotic fluctuations (Figure 3B), and thus we sought to characterize overall system dynamics as a function of environmental structure. We performed simulations with uncorrelated initial conditions across a range of pillar sizes and spacings, and classified system dynamics as ‘limit cycle’ or ‘chaotic’ by calculating the temporal autocorrelation of the spatial species distribution. If the spatiotemporal autocorrelation of all three species (minus steric barriers) at time *t* reached an autocorrelation above a threshold of 0.8 two or more times after *t*, we defined the dynamic state as cyclic at time *t* (see Supplemental Figure 8 & Methods). With this definition, we classified the dynamics as a function of *R* and Δ*x* into pseudo-phase diagrams for fraction of time spent in cyclic dynamics (Figure 5A) and the extinction frequency over the simulation time scale (Figure 5B). Example simulations are provided in Supplemental Movies 3-6. We found that smaller and more densely packed pillars lead to greater destabilization, with less time spent in limit cycle dynamics and higher rates of extinction. Intriguingly, however, with the smallest and most densely packed pillar structures we observed a reduced extinction frequency, reversing the trend seen at larger pillar spacings (Figure 5B, bottom row). This appears to be specific to the mechanisms by which pillars destabilize the system. With large pillars and spacings, spiral waves develop in open areas and are largely unperturbed by the pillars, resulting in cyclic behavior and few extinctions (Supplemental Movie 3). As pillar spacing decreases, open areas narrow to the point that spiral wave centers are destabilized, migrating erratically and eventually collapsing due to interference from other wave fronts (Supplemental Movie 4). With smaller pillar radii, the pillars themselves often act as wave centers, and appear to be particularly vulnerable to disruption via interference (Supplemental Movie 5). However, when small pillars are so densely packed that a pillar cannot serve as a wave center, the centers again migrate erratically between pillars, but the pillar density is high enough to ‘cage’ the rapidly diffusing wave centers and prolong their existence in a chaotically fluctuating state (Supplemental Movie 6). Thus, the prevalence of extinction cascades is a non-monotonic function of pillar density, suggesting that intermediate scales of spatial structure produce the strongest destabilizing effects on intransitive communities. Finally, to ensure that the observed changes to system dynamics and corresponding destabilizing effects were not dependent on the symmetry of a triangular lattice, we performed a subset of simulations where pillar radii or spacing were independently and randomly perturbed, and no significant changes to system dynamics and ecological outcomes were observed (Supplemental Figure 9).

**Figure 5:**
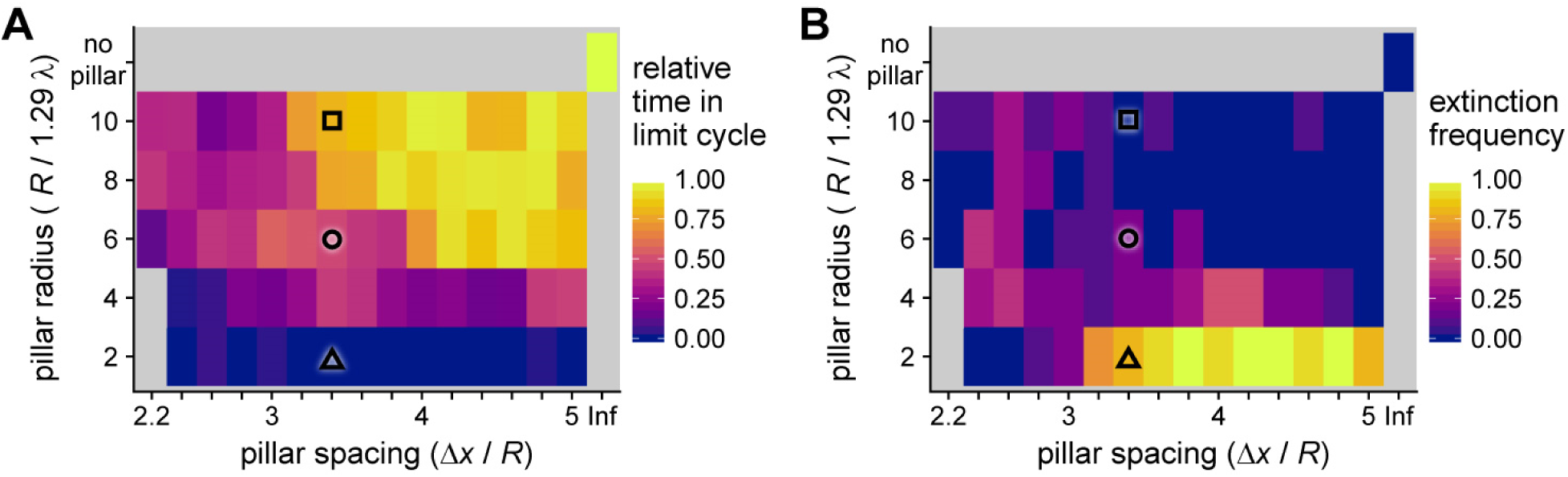
Severity of chaotic disruptions depend on structural characteristics of the environment. Over 1,000 doubling times, **A** shows the fraction of time the system displayed cyclic dynamics and B shows extinction frequency, both as functions of the size and spacing of the pillar array. 10 simulations were performed for each grid point. For each simulation, the amount of time spent in a limit cycle was normalized by the average time that isotropic simulations were classified as cyclic; this adjusts for systematically acyclic periods such as the grow-in phase. Three primary dynamic regimes were identified: (i) stable cycles, with larger and widely-spaced pillars; (ii) a transitory region at intermediate pillar size and spacing, where communities tend to either relax into a limit cycle or collapse; and (iii) when pillars are small and densely packed, unstable chaos with rapid community collapse. Simulation parameters are *L* / (1.29 *λ*) = 100 and *P* = 0.1, with pillar size and spacing as indicated in the figure. The simulations at *R* / (1.29 *λ*) = 2 and 4 with Δ*x* / *R* = 2.2 were omitted because the pillar spacing did not allow for accurate simulation of diffusion. Black symbols correspond to simulation conditions whose extinction time distributions were analyzed in Figure 6.

#### A three-state kinetic model describes coupling of dynamic transitions and extinction

In our three-species simulations we observed transitions from chaotic dynamics to limit cycles and back again, with many simulations ultimately making the transition from chaotic dynamics to the fully absorbing state of extinction. Though the simulations are deterministic, the ensemble of initial conditions create statistical variability in system dynamics. Thus, we wanted to characterize how the distribution of extinction times, and hence the time scale of coexistence, depended on environmental structure. We developed a three-state kinetic model to describe transitions between chaotic (C), limit cycle (L), and extinct (E) states, using three positive rate parameters to connect the states *(k_CL_, k_LC_*, and *k_CE_)*. The closed-form solution to our model (Supplemental Text 2) predicts that all systems with structural perturbations will go extinct in the infinite time limit, which is consistent with previous work (18, 19). It also predicts that the rates of arrival to the extinct state depend on the dynamics fostered by the environmental structure. To test this, we used structural conditions whose initial dynamics were classified as either limit cycle, chaotic, or mixed for the first 1,000 doubling times (marked tiles in Figure 5), and fit the observed distribution of arrival times as a function of environmental structure to those predicted by the model over a period of 10,000 doubling times. We found that our model recapitulated observed distributions of extinction times (Figure 6), and that indeed, changes in environmental structure had significant effects on the distribution of extinction times. These results indicate that structurally-induced destabilization results from a combination of decreased rates of transition from chaotic fluctuations to limit cycle dynamics and/or increased rates of transition from chaotic dynamics to extinction (see model diagrams in Figure 6). Accordingly, systems that remained largely in a limit cycle had slower rates of extinction. The fitted model parameters were functions of multiple individual transition rates with complex mappings (Supplemental Text 2), hence direct inference of the effects of structural perturbations on individual transition rates (e.g. from limit cycle to chaos) were not possible with this model.

**Figure 6:**
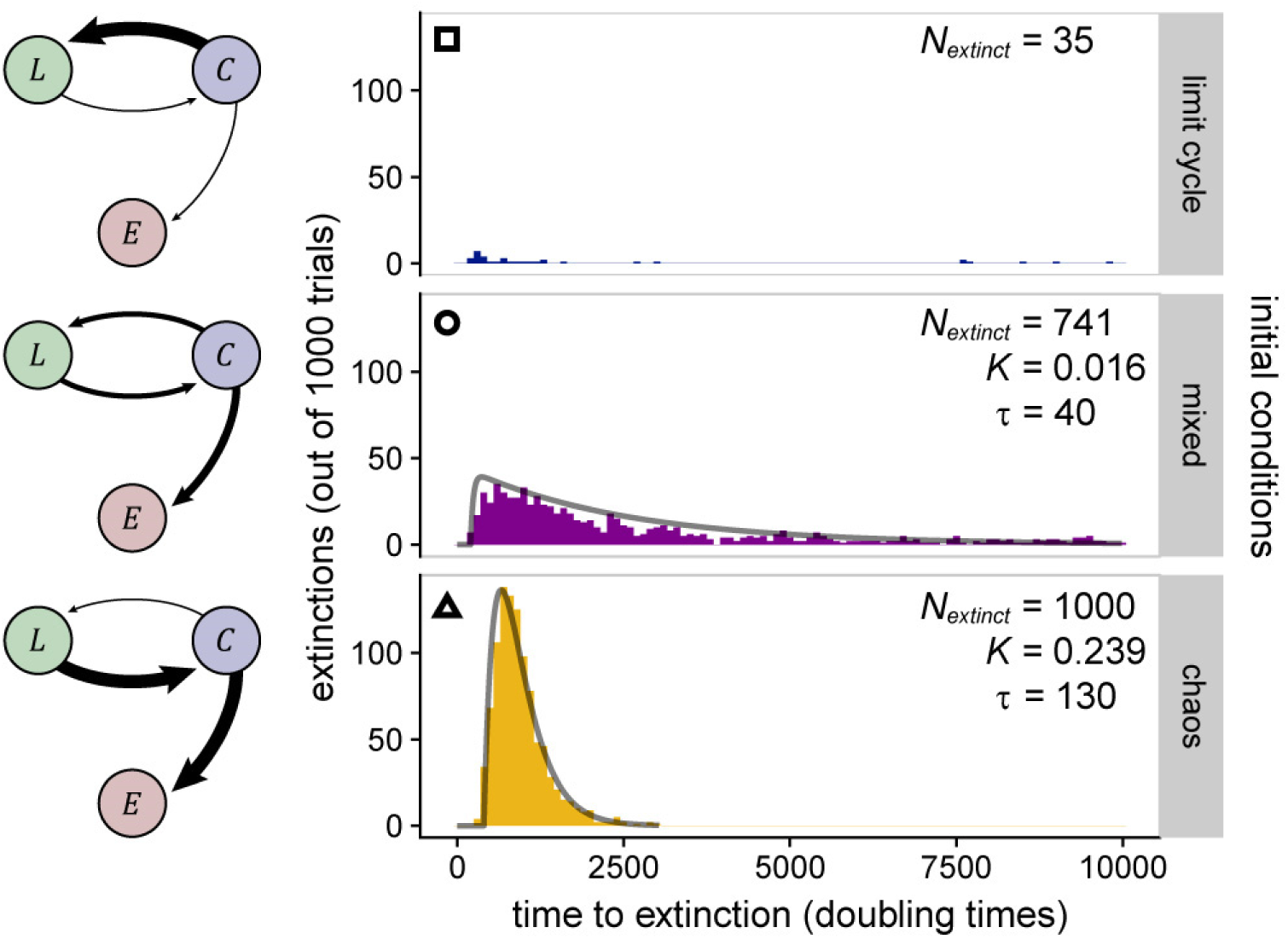
A kinetic model of dynamic transitions predicts extinction time distributions for a range of environmental structures. From the lattice structures indicated by overlaid symbols in Figure 5 (indicated here at top left in histogram plots), we performed 1,000 replicate simulations for 10,000 doubling times to measure the distribution of extinction times and compare them to our model predictions. These conditions typify the three observed dynamic regimes (limit cycle, transitory and chaotic), and map to a three-state model of system dynamics with two correlated rate parameters that depend on structural characteristics (Supplemental Text 2). The histograms were constructed from observed extinction times, and grey lines are fits to probability distributions predicted from the three-state model. Fitting was not attempted for the cyclic case (top row), as only 3.5% of simulations were observed to go extinct over the simulation period. The number of extinctions and (where applicable) the fit parameters are shown within the corresponding plots. At left, connections between the dynamical states of limit cycle (L), chaotic (C), and extinction (E) are depicted with relative rates qualitatively indicated by the width of the arrows. Simulation parameters are *L* / (1.29 *λ*) = 100 and *P* = 0.1, with pillar size and spacing as indicated in the figure.

#### Larger systems prolong species coexistence despite chaotic fluctuations

Lastly, we sought to characterize the effect of system size on community stability. Holding the structure of the pillar array constant, we observed that the mean time to an extinction cascade increased approximately exponentially with increasing system size (Figure 7A). This suggests that with sufficiently large systems relative to the natural length scale, communities can coexist for long periods despite continual chaotic fluctuations in individual species abundances and distributions. However, consistent with the predictions of our kinetic model (Figure 6), larger systems cannot fully prevent extinctions, as evidenced by observed extinction frequencies when simulation times were extended. In Figure 7B we show that for a given simulation duration there is a system size above which the extinction frequency drops to nearly zero, however, simply extending the simulation time can push the extinction frequency to unity.

**Figure 7:**
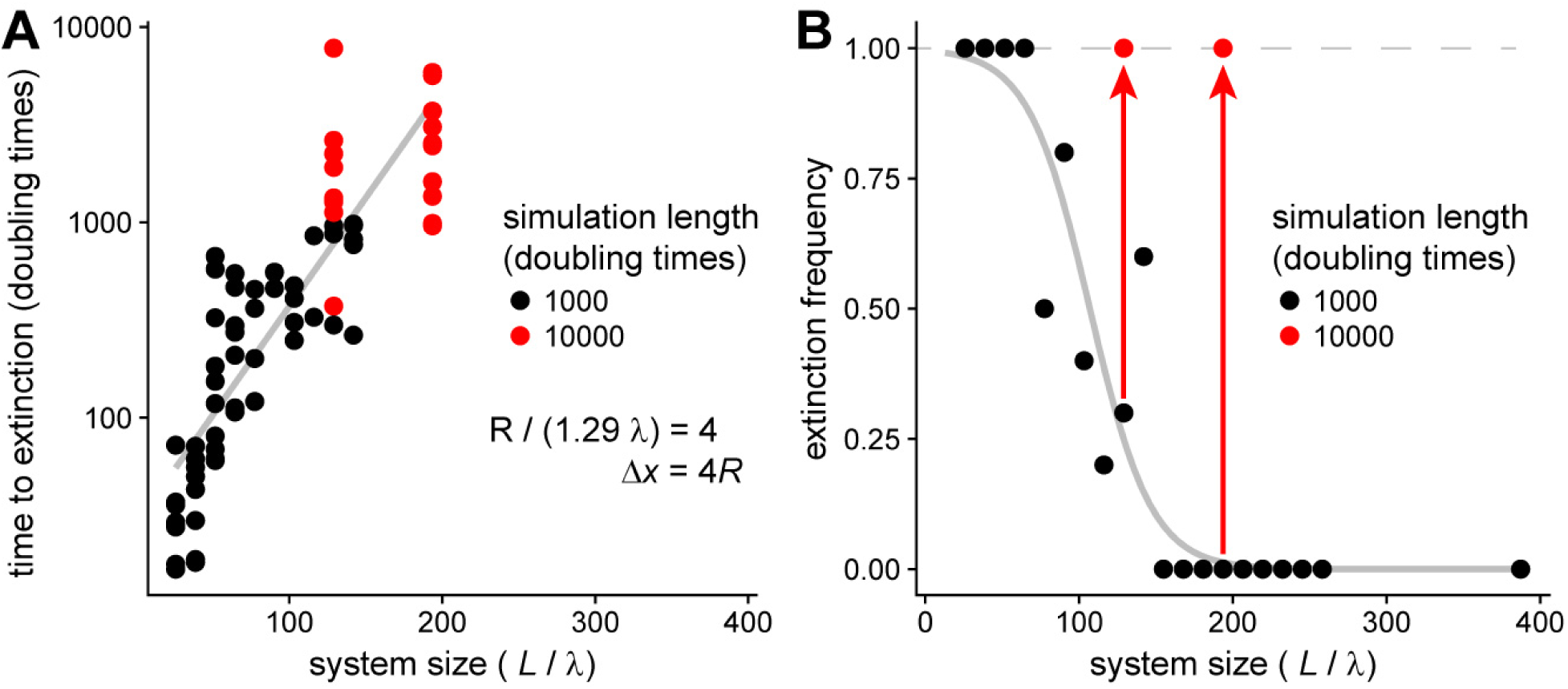
Larger system sizes delay but do not prevent extinction cascades. **A**, plotting time to extinction as a function of system size reveals an approximately exponential relationship, suggesting that large systems can persist in a state of chaotic fluctuation for long time periods. Each point is a single simulation, with 10 replicate simulations per system size *L*. **B**, extinction frequency as a function of system size. The black points and fitted grey logistic regression curve suggest a critical system size at which extinctions are no longer observed for the fixed number of simulated doubling times (here, 1,000). However, by increasing simulation duration, the observed extinction frequency saturates to approximately 1 (red points and arrows), indicating that large systems delay but not prevent extinction, consistent with our model which predicts that all anisotropic environments will eventually end with an extinction cascade. Simulation parameters are *R* / (1.29 *λ*) = 4, Δ*x* = 4 *R*, and *P* = 0.1, with system size *L* as indicated in the figure.

## DISCUSSION

Using *in silico* simulations of ecological communities, we found that addition of structural complexity to the environment results in fundamental changes to community dynamics and outcomes in a manner dependent on the specific interaction network topology. Specifically, we observed that for two mutually competitive species, structurally complex environments allowed for long-term coexistence between species with relatively large differences in competitive fitness, an outcome impossible in well-mixed or isotropic environments. Conversely, for a three-species intransitively competing community, which is expected to be stable under isotropic conditions (5), we found that environmental structure can disrupt the dynamic spatial patterns that stabilize these communities, resulting in chaotic fluctuations in species abundances and spatial distributions, and an increased frequency of extinction cascades. Together, these findings strongly suggest that the physical structure of the environment can interact significantly with the specific nature of interspecies interactions within resident communities to affect stability and dynamics, and more generally indicate that physical attributes of the environment must be considered when assessing the stability of resident communities.

Our results extend established findings that spatially structured communities maintain biodiversity by localizing interactions among community members (7, 21, 22). In particular, in the context of simple competition the spatial bottlenecks that structurally complex environments provide impede competitive mechanisms to the point that only a small fraction of a given population is engaged in active competition, and hence fitness differences become less important relative to geometric advantages provided by specific localization within the environment. However, our findings also suggest that intransitive interaction networks are not a robust means of stabilizing communities, as has been theoretically postulated (23, 24). Likewise, if deviation from isotropic conditions (which is found in virtually all natural environments) only serves to accelerate the frequency of extinction cascades within these networks, this work offers a mechanism as to why such networks are only rarely observed outside of the lab (25-27). We speculate, based on scaling effects, that the increase in surface area-to-volume ratio going from 2D into 3D will only enhance the stabilization of asymmetric competition between two species. Conversely, given the potential augmentation of structural complexity available in higher dimensions, we expect that under similar conditions chaotic fluctuations would be a robust feature of intransitively competing communities. We also expect that the shape of the steric barriers will play a non-trivial role in ecosystem dynamics and stability; we chose circles for simplicity, as they are characterized by a single parameter. The spectrum of available interface curvatures within a particular environmental structure is a function of both overall spatial scale (e.g. here Δx), and the shape of the steric objects themselves. Rationally designed structures could be used to tune the range of competitive asymmetries and/or stochastic fluctuations that an environment can stably support, and to shift system dynamics and stability to favor particular interaction topologies. It is of interest to assess whether our findings are robust when placed in the context of other physical and ecological phenomena. For example, how robust are pinned competition interfaces to stochastic spatial fluctuations caused either by finite organism size or other forms of motility (besides diffusion), tunable interaction strengths, such as with competition sensing (28, 29), or phenotypic differentiation (30)? Are chaotic fluctuations a dominant dynamic state when cells can respond to chemical gradients via chemotaxis? What are the effects of physical structure on species distributions for larger networks, where specific interaction motifs are embedded within a more complex ecological context? These extensions will pave the way toward future theoretical work, as well as generating specific hypotheses to be tested experimentally.

Finally, we note that the reductionist approach we take here is valuable toward unravelling the multitude of forces acting on microbial communities in complex environments. While we focus specifically on environmental structure, and others give similar focus to flow (31, 32) and chemical gradients (33) in structuring communities, all of these environmental features are intimately linked and in combination will modulate impacts on communities in important ways (9). Building a bottom-up understanding of how various features interact to drive community processes is therefore essential in determining the primary forces acting on a community in a given environmental context, paving the way toward the ultimate goals of understanding basal mechanisms of ecosystem dynamics and of targeted and robust interventions in microbial communities.

## METHODS

### Two species mutual killer simulations

Simulations were randomly seeded with pink noise (34) at an average density of 10% of the carrying capacity, with each species represented by its own field matrix. Pillars were placed in a triangular lattice with the specified radius and spacing. Microbial density that coincided with pillar locations was removed from the simulation. The bounding box and pillar edges were modeled as reflecting boundary conditions. At each simulated time step (Δ*t* = 0.1*t*, with *t* in doubling times), populations diffused via a symmetric Gaussian convolution filter with standard deviation set by the diffusion coefficient, 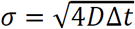. After the diffusion step, changes in population density (growth and death) were calculated using the equations given in the main text, and used to update the density of each species. Hard upper and lower bounds (1 and 0.001 in units of carrying capacity, respectively) were enforced to improve numerical stability of simulations; populations densities outside this range were set to 1 and 0, respectively. For each set of lattice constants and competitive asymmetry values, 30 independently initialized replicates were simulated for 2000 doubling times. Mean population abundances and images of the simulation were recorded at an interval of 0.4*t* for the duration of the simulation. Extinction was defined as the mean population density of either species dropping below a threshold value of ((2*R)*^2^ − π*R^2^)*/4*A*, where *R* is the pillar radius and *A* the area of lattice points not obstructed by pillars, to account for surviving populations ‘trapped’ between a pillar and the corner of the simulation box and therefore not in contact with the rest of the simulation.

### Calculation of pinned curvature

To obtain higher resolution of pinned curvature in asymmetric competition, two pillars of *R* = 12.9 *λ* were put at two opposing edges of a simulation box, and in contact with the simulation boundary leaving a single gap between the pillars. Two competing species were symmetrically and uniformly inoculated at 30% of the carrying capacity on either side of this gap, leaving a single flat interface spanning the distance between the two pillars. Simulations were then allowed to evolve as above until dynamics ceased due to either pinning or extinction. All combinations of the indicated competitive asymmetries were sampled, and pillar gap distances were sampled by varying the size of the simulation box. For simulations where pinning was observed, the interface location was defined as the boundary points where species A and B were of equal abundance. The interface curvature was calculated from three points along that boundary (the midpoint and the two points in contact with the pillars); this method was found to be more robust than other circle-fitting methods, especially for low curvatures and narrow pillar gaps.

### Intransitive three-species simulations

Three species intransitive simulations were carried out similarly to the two-species cases described above, with the competition terms in the model modified to reflect the intransitive interaction network topology. Simulations were inoculated randomly with 10 replicate simulations per structural condition. Unless otherwise indicated, simulations were evolved for 1,000 doubling times, with images written every 0.2t.

A schematic of the classification of simulation dynamics is given in Supplemental Figure 1. Spatiotemporal autocorrelations were calculated for each simulation, where correlations at each time point were calculated from the concatenated vectorized simulation matrix of all non-pillar grid locations for each species, i.e. for the autocorrelation matrix in Supplemental Figure 1, each matrix entry represents the correlation of two 438,000 (3 species multiplied by 146,000 unique non-pillar grid locations) length vectors at the indicated time points. Using the autocorrelation matrix, at every time point (i.e. starting from the matrix diagonal and moving forward in time), that time point was classified as exhibiting limit cycle dynamics if the autocorrelation rose above the threshold value of 0.8 for at least two cycles. This threshold was chosen empirically as the level at which isotropic simulations were reliably classified as limit cycles over the duration of the simulation (excluding grow-in periods and final time points for which future dynamics were not observed). Extinction events were calculated as in the two-species cases.

To establish correlated initial conditions (Figure 4), the following procedure was used: for each replicate set of simulations, a random initial inoculum at density 10% of the carrying capacity was generated using the same random seed (i.e. constructing 10 identical initial condition matrices). Then, for each individual replicate, a randomly selected percentage (as indicated in Figure 4) of non-pillar grid locations were randomly resampled between 0 and 10% of the carrying capacity. Simulations were then allowed to evolve as described above. At each time point, each unique pairwise correlation (45 for the 10 replicates used) between vectorized simulation matrices was calculated, and the mean over all pairwise correlations was used to generate Figure 4. Correlation traces were truncated upon the first observed extinction event among the replicates.

### Kinetic modeling

For details on assumptions and analysis of the kinetic state model, and derivation of the closed-form solutions for the time-to-extinction distributions, see Supplemental Text 2. Histograms were generated from randomly initialized simulations as described above, with 1,000 replicates per set of lattice constants, each over 10,000 doubling times. Model parameters *K* and *τ* were fit by minimizing the squared error between the empirical cumulative distribution function (CDF) from simulated data and the corresponding CDF predicted by the model; global minima in the parameter space were found using grid search. A temporal offset parameter *τ*_offset_ was also fit to account for grow-in periods, effectively shifting the histogram along the time axis and setting extinction probability for *t < τ*_offset_ to zero.

## Data availability

Code to run simulations and analyses, as well as processed data from raw images will be posted on Github.

## ACKNOWLEDGEMENTS

We thank Kerwyn Huang, Will Ratcliff, Kalin Vetsigian, Ajay Gopinathan, Raghu Parthasarathy, and Brendan Bohannan for helpful discussions and comments on the manuscript. We also thank Rob Yelle and Michael Coleman for assistance with high performance computing resources at the University of Oregon. Research reported in this publication was supported in part by the National Institute of General Medical Sciences of the National Institutes of Health under award number 1P50GM098911, and the University of Oregon to TSU. This work was performed in part at Aspen Center for Physics, which is supported by National Science Foundation grant PHY-1607611. The content is solely the responsibility of the authors and does not necessarily represent the official views of the National Institutes of Health.

## SUPPLEMENTAL MOVIE LEGENDS

**Supplemental Movie 1: Symmetric two-species competition in structurally isotropic and anisotropic environments.** Simulation parameters are *L* / (1.29 *λ*) = 100, *P* = 0.1, with *R* / (1.29 *λ*) = 2 and Δ*x* = 3.4 *R* for the anisotropic case. The movie depicts system dynamics over 360 doubling times; the anisotropic simulation is pinned after approximately 60 doubling times.

**Supplemental Movie 2: Pinning and coexistence of species with asymmetric competitive fitness.** Simulation parameters are *L* / (1.29 *λ*) = 100, *P_A_* = 0.112, *P_B_* = 0.088, *R* / (1.29 *λ*) = 2 and Δ*x* = 3 *R*. The movie depicts system dynamics over 85 doubling times.

**Supplemental Movie 3: Large and widely spaces pillars do not significantly perturb intransitive communities.** Simulation parameters are *L* / (1.29 *λ*) = 100, *P* = 0.1, *R* / (1.29 *λ*) = 8, Δ*x* = 5 *R*. The movie depicts system dynamics over 1,000 doubling times.

**Supplemental Movie 4: Dense pillars induce wave destabilization and community collapse.** Simulation parameters are *L* / (1.29 *λ*) = 100, *P* = 0.1, *R* / (1.29 *λ*) = 10, Δ*x* = 2.6 *R*. The movie depicts system dynamics over 1,000 doubling times.

**Supplemental Movie 5: Pillars may serve as unstable wave centers.** Simulation parameters are *L* / (1.29 *λ*) = 100, *P* = 0.1, *R* / (1.29 *λ*) = 6, Δ*x* = 3.4 *R*. The movie depicts system dynamics over 1,000 doubling times.

**Supplemental Movie 6: Small dense pillars can ‘cage’ wave centers and prolong community coexistence under chaotic dynamics.** Simulation parameters are *L* / (1.29 *λ*) = 100, *P* = 0.1, *R* / (1.29 *λ*) = 2, Δ*x* = 2.4 *R*. The movie depicts system dynamics over 1,000 doubling times.

